# Artificial light at night amplifies seasonal relapse of haemosporidian parasites in a widespread songbird

**DOI:** 10.1101/2020.06.22.163998

**Authors:** Daniel J. Becker, Devraj Singh, Qiuyun Pan, Jesse D. Montoure, Katherine M. Talbott, Sarah Wanamaker, Ellen D. Ketterson

## Abstract

Urban habitats can shape interactions between hosts and parasites by altering not only exposure rates but also within-host processes. Artificial light at night is common in urban environments, and chronic exposure can impair host immunity in ways that may increase infection. However, studies of causal links between this stressor, immunity, and infection dynamics are rare, particularly in migratory animals. Here, we experimentally tested how artificial light at night affects cellular immunity and haemosporidian parasite intensity across the annual cycle of migrant and resident subspecies of the dark-eyed junco (*Junco hyemalis*). We monitored an experimental group exposed to light at night and a control group under natural light/dark cycles as they passed through short days simulating early spring to longer days simulating the breeding season, followed by fall migration. Using generalized additive models, we show that artificial light at night increased inflammation, and leukocyte counts were greatest in early spring and fall. At the start of the experiment, few birds had active infections based on microscopy, but PCR revealed many birds had chronic infections. Artificial light at night increased parasitemia across the annual cycle, with strong peaks in spring and fall that were largely absent in control birds. As birds were kept in indoor aviaries to prevent vector exposure, this increased parasitemia indicates relapse of chronic infection during costly life history stages (i.e., reproduction). Although the immunological and parasitological time series were in phase for control birds, cross-correlation analyses also revealed artificial light at night desynchronized leukocyte profiles and parasitemia, which could suggest a general exaggerated inflammatory response. Our study shows how a common anthropogenic influence can shape within-host processes to affect infection dynamics.

## Introduction

As suburban and urban habitats expand globally, certain species are attracted to these environments owing to a combination of altered microclimate, predation risk, and resource availability [1,2]. Such changes in biotic and abiotic conditions can have cascading and complex effects on wildlife [3,4]. When considering infectious disease, urban habitats can shape the interactions between hosts and parasites by altering within-host processes such as resistance or tolerance [5]. Understanding how urbanization affects these processes is especially important, as it can facilitate novel cross-species transmission opportunities between various host species [6].

One aspect of urban habitats that could have strong effects on these within-host processes is artificial light at night (ALAN). By altering animal circadian rhythms, ALAN can cause a suite of physiological changes relevant to infection [7]. In particular, chronic exposure to realistic urban levels of ALAN (e.g., 5 lux) can suppress host immune function, including cellular response and microbial killing ability [8,9]. Prolonged ambient light exposure at similar or greater intensities (e.g., 5 to 400 lux) can also dysregulate the inflammatory response, causing inflammation with pathological effects [10,11]. Ultimately, these changes likely alter host infection outcomes and parasite transmission [12]. In a rare test of causal relationships, similar levels of ALAN (i.e., 8 lux) altered gene regulatory networks and extended the infectious period for West Nile virus in house sparrows (*Passer domesticus*) [13]. This suggests that ecologically relevant exposure to ALAN could enhance the transmission of parasites to arthropod vectors and thus increase epidemic potential; however, the consequences of ALAN on infectious disease dynamics in wild hosts generally remain poorly understood.

Although the impacts of ALAN on immunity have been studied in various captive and free-living animals, these consequences have been less explored in migratory species. For birds in particular, many species migrate at night [14], facilitating exposure to light pollution along major migration routes [15]. Many birds also use suburban and urban habitats as stopover sites [16], and widespread ALAN in these environments can further affect habitat selection during migration [17,18]. Because physiological preparation for long-distance migration can itself suppress immune function [19,20], any effects of ALAN on within-host processes could be especially pronounced in migrants. If such changes amplify parasite transmission, urban habitats could become ecological traps for migratory species and in turn hotspots of spillover risk [21].

Here, we experimentally tested how ALAN exposure affects immunity and infection across the annual cycle of migratory and resident dark-eyed juncos (*Junco hyemalis*). Juncos are broadly distributed across North America [22], composed of subspecies that diverged 15,000 years ago and vary in migratory behavior, reproductive timing, and morphology [23]. Migratory slate-colored juncos (*J. h. hyemalis*) breed across Canada and the northern United States, migrating south in autumn. In the Appalachian Mountains, resident juncos (*J. h. carolinensis*) are sympatric with overwintering migrants from fall through early spring, when *J. h. hyemalis* initiate northern migration and *J. h. carolinensis* transition into breeding condition. To recreate the annual cycle of residents and migrants captured during sympatry, we modified photoperiod, which provides a consistent and predictive cue for animals to coordinate seasonal events such as reproduction and migration [24]. Mirroring natural changes in daylength across six months allowed us to capture these key life history events from early spring through late fall.

To determine effects of ALAN on within-host infection processes across these annual cycles in migrants and residents, we quantified cellular immunity and parasitemia. We focused on haemosporidians (i.e., causative agents of avian malaria) [25], which are common in juncos and are transmitted by arthropod vectors from spring through autumn [26,27]. Haemosporidians cause a short, acute infection followed by a chronic phase characterized by low parasitemia and latency in peripheral host tissue [25,28]. Parasites present in hosts sampled in winter therefore signal chronic infections [27,29]. However, seasonal relapse of chronic infections commonly occurs from physiological changes in preparation for bird reproduction or migration [29–31]. Manipulating photoperiod in an indoor aviary free of potential vectors allowed us to isolate effects of ALAN and seasonality on within-host processes, whereas field comparisons may also have greater exposure to vectors in urban habitats [32]. We predicted ALAN would promote inflammation and hematological indices of physiological stress (i.e., ratios of heterophils to lymphocytes [33]). Under control conditions, we expected these cellular immune profiles and parasitism to generally be synchronized; however, immune dysregulation from ALAN, as reflected by desynchronization (i.e., a phase lag) of these relationships, could cause chronic infections to reactivate. Because stages of the annual cycle carry distinct energetic costs, we also expected these relationships would vary by photoperiod and migratory strategy. Parasitemia could be greatest under ALAN in late spring, when migrants prepare for northern migration and residents start summer breeding. However, migrants could display greater parasitemia than residents in early autumn, when residents complete breeding while migrants initiate their southern migration.

## Methods

### Junco sampling and housing

In December 2017, we captured adult male migrant and resident juncos at the Mountain Lake Biological Station (Pembroke, VA). Each bird was fitted with an aluminum U.S. Fish & Wildlife Service leg band. Subspecies were differentiated by bill color, plumage, and size [22]. Residents (*n* = 18) and migrants (*n* = 24) were housed with *ad libitum* food and water in an indoor aviary at Indiana University in Bloomington, Indiana. During a prior experiment, we gradually increased then decreased photoperiod to simulate a full annual cycle through 2018 [34], and birds were maintained in a photosensitive state at 9 hours daylength (i.e., 9L) until our experiment in 2019.

### Experimental design

We randomly housed equal numbers of residents and migrants in mixed-flock, free-flying rooms in two separate treatments. Per treatment, birds were held in two interconnected and equal-sized rooms (2.59 by 2.06 meters). To minimize room artifacts, all rooms were maintained at equal temperature (18 ± 2°C), were on the same side of the aviary, had the same door and window placement, and had equal numbers of perches and food and water dishes. Starting in February 2019, we simulated another full annual photoperiodic cycle while exposing control birds to natural light/dark (LD) cycles and treatment birds to constant ~2.5 ± 0.5 lux average intensity during night (ALAN). ALAN intensity was determined from a dose-dependent study in great tits, in which 0.5–5 lux increased reproductive activity [35], and such levels reflect ALAN exposure of urban birds [36]. All rooms blocked natural sunlight and provided photoperiod light with floor lamps via fluorescent bulbs (Phillips, 32W; full spectrum white light). Daylight was maintained at 140 ± 5 lux average intensity, and LD birds were maintained at total darkness at night.

We increased photoperiod every fourteen days by 0.3–1 hour from February 16 to September 2 (9L:15D to 16L:8D), continued for eight weeks at 16L:8D, and then decreased photoperiod until November 22 (16L:8D to 9L:15D). At each timepoint, we captured free-flying indoor birds 30 minutes after the onset of morning photoperiod light. For a complementary study of how ALAN affects reproductive physiology, birds received a standardized intramuscular challenge of gonadotropin releasing hormone at each timepoint [37]. Thirty minutes after injection, we collected 50–100 μL blood from the brachial vein using sterile 26G needles and made thin blood smears for hematology. Birds were kept in opaque bags between injections to reduce stress, and the 30 minutes of holding time between challenge and bleeding was uniform across birds and is unlikely to affect circulating leukocyte counts [33]. We collected repeated blood smears across nine timepoints (Table 1), which we chose from photoperiodic thresholds in junco reproductive condition and timing of the junco annual cycle [22,34]. Residents and migrants initiate elevation of testosterone at 11L and 12L, respectively, followed by gonadal recrudescence at 12.4L and 13L. These photoperiods correspond to early spring daylengths and were followed by increasing daylengths simulating the breeding season, for which maintenance at 16L promotes a photorefractory state to terminate reproduction. Declining daylengths end this refractory state and simulate onset of fall migration to complete a full annual reproductive cycle. Some birds were removed from the study due to injury or mortality, resulting in 21 migrants (LD=10, ALAN=11) and 12 residents (LD=5, ALAN=7) repeatedly sampled for analyses.

**Table 1.** Experimental sampling timepoints and corresponding photoperiod.

| Week | Photoperiod |
| --- | --- |
| 3/18/2019 | 10L:14D |
| 4/15/2019 | 11.4L:12.6D |
| 5/27/2019 | 12.5L:11.5D |
| 7/8/2019 | 15L:9D |
| 9/2/2019 | 16L:8D |
| 10/10/2019 | 12L:12D |
| 10/24/2019 | 11L:13D |
| 11/7/2019 | 10L:14D |
| 11/21/2019 | 9L:15D |

### Hematology and PCR

Blood smears were stained with buffered Wright–Geimsa (Astral Diagnostics Quick III). From each sample (*n =* 297), we evaluated leukocyte profiles and parasite intensity using light microscopy (AmScope, B120C-E1). A single observer, blind to the treatment and migratory strategy of each bird, first recorded the number of white blood cells under 40X magnification across 10 random fields to estimate inflammatory state and investment in cellular immunity. Next, a differential count was performed by recording the identity of the first 100 leukocytes at 1000X magnification [38]. Approximately 100 fields of view at 1000X magnification were screened for *Plasmodium, Haemoproteus,* and *Leucocytozoon* [25]. We derived the mean total white blood cell count (total leukocytes), ratio of heterophils to lymphocytes (HL ratios), and total number of haemosporidian parasites (parasite intensity) as three separate response variables.

As microscopy primarily detects active haemosporidian infection [25,28], we used PCR to detect chronic infections at the start of our experiment (i.e., 9L). We extracted genomic DNA from erythrocytes in Longmire’s solution using RSC Whole Blood DNA Kits (Promega) run on a Maxwell RSC at Indiana University, and DNA was eluted in 70 μL buffer. We used previously published primers and PCR protocols to amplify a conserved fragment of the haemosporidian mitochondrial rRNA gene common to *Plasmodium, Haemoproteus,* and *Leucocytozoon* [39,40]. We re-screened any negative birds after initial PCR to reduce possible risks of false negatives.

### Statistical analysis

We analyzed all response variables using R [41]. We first used generalized additive mixed models (GAMMs) with the *mgcv* package to account for potential nonlinear effects of seasonality (i.e., photoperiod) and treatment [42]. We analyzed data from migrants and residents separately, as initial analyses suggested no difference in any of these three outcomes (i.e., total leukocytes, HL ratios, parasitemia) by migratory strategy or its interaction with treatment (Table S1). For each response variable per migrants and residents, we fit a GAMM with treatment, a smoothed effect of photoperiod, and their interaction using thin plate splines [43]. GAMMs included a random effect of individual band number to account for repeated measures. We modeled both total leukocytes and HL ratios with Tweedie distributions, whereas we used a zero-inflated Poisson distribution for parasitemia.

To test temporal dependence between cellular immunity and infection, we estimated the cross-correlation between log-transformed parasitemia and (*i*) log-transformed total leukocytes and (*ii*) square-root HL ratios for all photoperiod time lags between each time series [44]. We estimated cross-correlation coefficients for each treatment within migrants and residents. We then fit an additional set of GAMs with a smoothed effect of photoperiod lag, treatment, and their interaction with cyclic cubic splines and a Gaussian distribution for each strata of the data. Strong, positive correlations around the zero lag would support synchrony between the time series (i.e., in phase), whereas weaker correlations across broader photoperiod lags would suggest desynchronization of immunity and parasitemia and possibly immune dysregulation.

## Results

### ALAN effects on leukocyte profiles

ALAN increased total leukocytes compared to LD birds (Fig. 1A), and effect sizes were similar for residents and migrants (Table 2). Regardless of migratory strategy, we also found strong seasonal (i.e., photoperiodic) variation in total leukocytes, with greater inflammation at the start and end of our experiment (i.e., early spring and late fall). Although GAMMs explained up to 59% of the deviance in total leukocytes, seasonal patterns did not vary by treatment. In contrast, we found weaker effects of treatment and photoperiod on HL ratios (Table S2). ALAN did not broadly affect HL ratios (Fig. 1B). Only migrants showed an effect of photoperiod on HL ratios, for which this measure of physiological stress was highest in ALAN birds during early spring.

**Figure 1.**
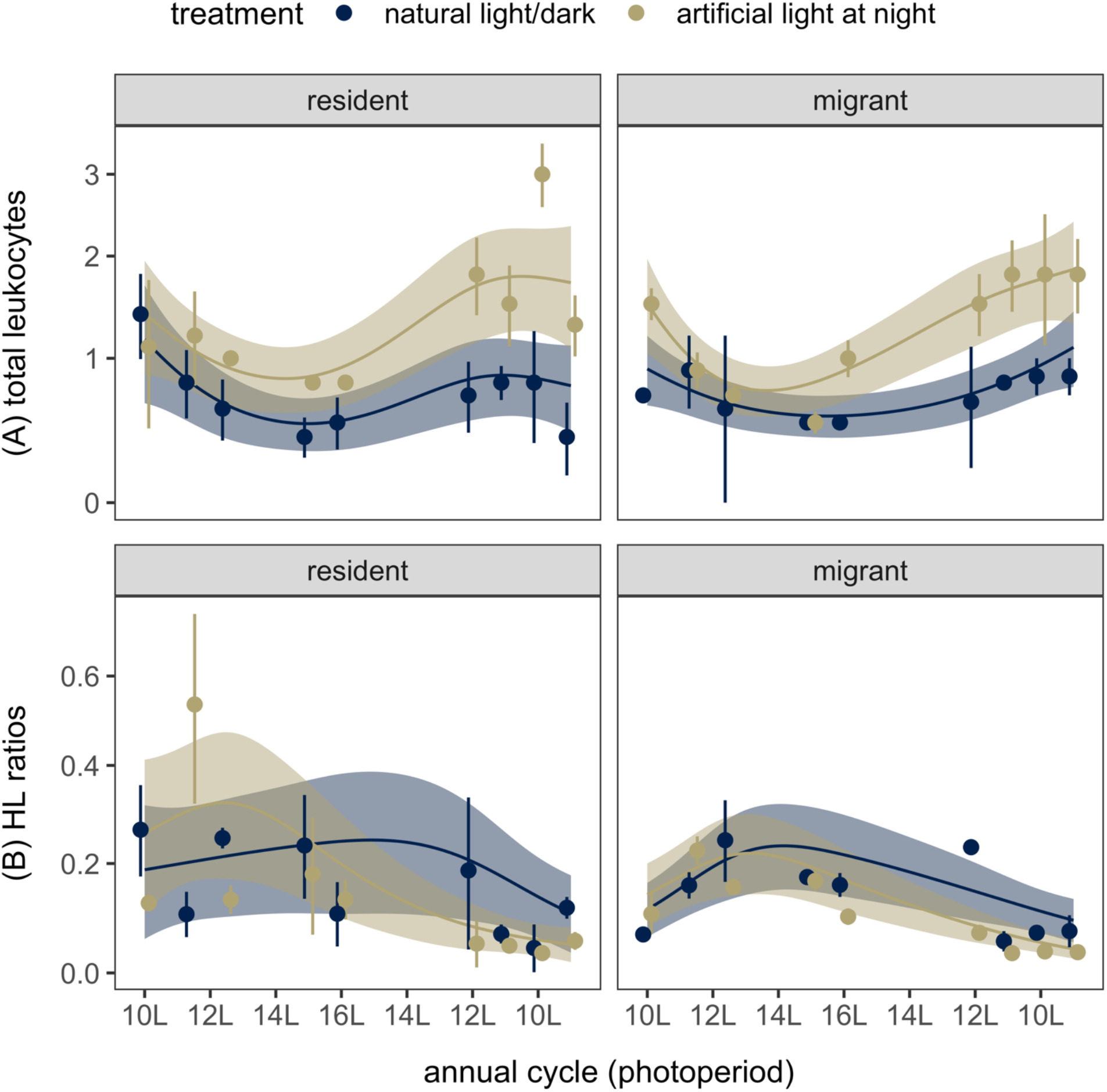
Effects of ALAN on (A) total leukocytes and (B) HL ratios across the simulated annual cycle (i.e., photoperiod, L) of resident and migrant dark-eyed juncos. Points show medians with jackknife estimates of standard error and are jittered to reduce overlap. Lines and shaded bands show the fitted values and 95% confidence interval from each GAMM. The vertical axes are displayed with a modulus transformation to accommodate lower bounds at zero.

**Table 2.** Effects of ALAN treatment, photoperiod, and their interaction on total leukocytes. GAMMs were stratified by migratory strategy and fit using restricted maximum likelihood. Fixed effects are presented with model coefficients (categorical) or the estimated degrees of freedom (EDF) and test statistics. Deviance explained (%) is presented for both GAMMs.

| Term | Residents<br>(deviance explained=52%) |  |  |  |  | Migrants<br>(deviance explained=59%) |  |  |  |  |
| --- | --- | --- | --- | --- | --- | --- | --- | --- | --- | --- |
| | $\beta$ | $t$ | EDF | $F$ | $p$ | $\beta$ | $t$ | EDF | $F$ | $p$ |
| Intercept | -0.30 | -1.88 |  |  | 0.06 | -0.28 | -2.51 |  |  | 0.01 |
| ALAN | 0.55 | 2.67 |  |  | 0.01 | 0.50 | 3.18 |  |  | <0.01 |
| s(photoperiod) |  |  | 3.27 | 5.32 | <0.01 |  |  | 2.90 | 7.03 | <0.01 |
| s(photoperiod, LD) |  |  | 0.50 | 0.34 | 0.60 |  |  | 0.00 | 0.07 | 0.99 |
| s(photoperiod, ALAN) |  |  | 1.00 | 2.34 | 0.13 |  |  | 3.13 | 2.96 | 0.07 |

### Haemosporidian intensity under ALAN

At our first blood smear collection (10L), the prevalence of active infection (i.e., infected erythrocytes detectable with microscopy) was 13.3% for LD birds (*n* = 15) and 16.7% for ALAN birds (*n* = 18); only ALAN residents had no microscopy-detectable infection (Fig. 2A). PCR confirmed that 53.3% of LD birds and 66.7% of ALAN birds had chronic infections at the start of the study (i.e., 9L; Fig. S1), and the odds of PCR-detectable infection did not differ by treatment or between migrants and residents (Table S3). Most PCR-negative birds at 9L did not show microscopically detectable infection at our first blood smear collection (10L, 92%; Fig. S2). Diagnostic mismatches suggest PCR-negative birds likely harbored latent infections (i.e., those present in peripheral tissue) that relapsed later in the experiment, although false negatives are also possible [25,45]. However, by the end of the study, active infections were common in all experimental groups, ranging from 30% to 100% prevalence (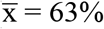; Fig. 2A).

**Figure 2.**
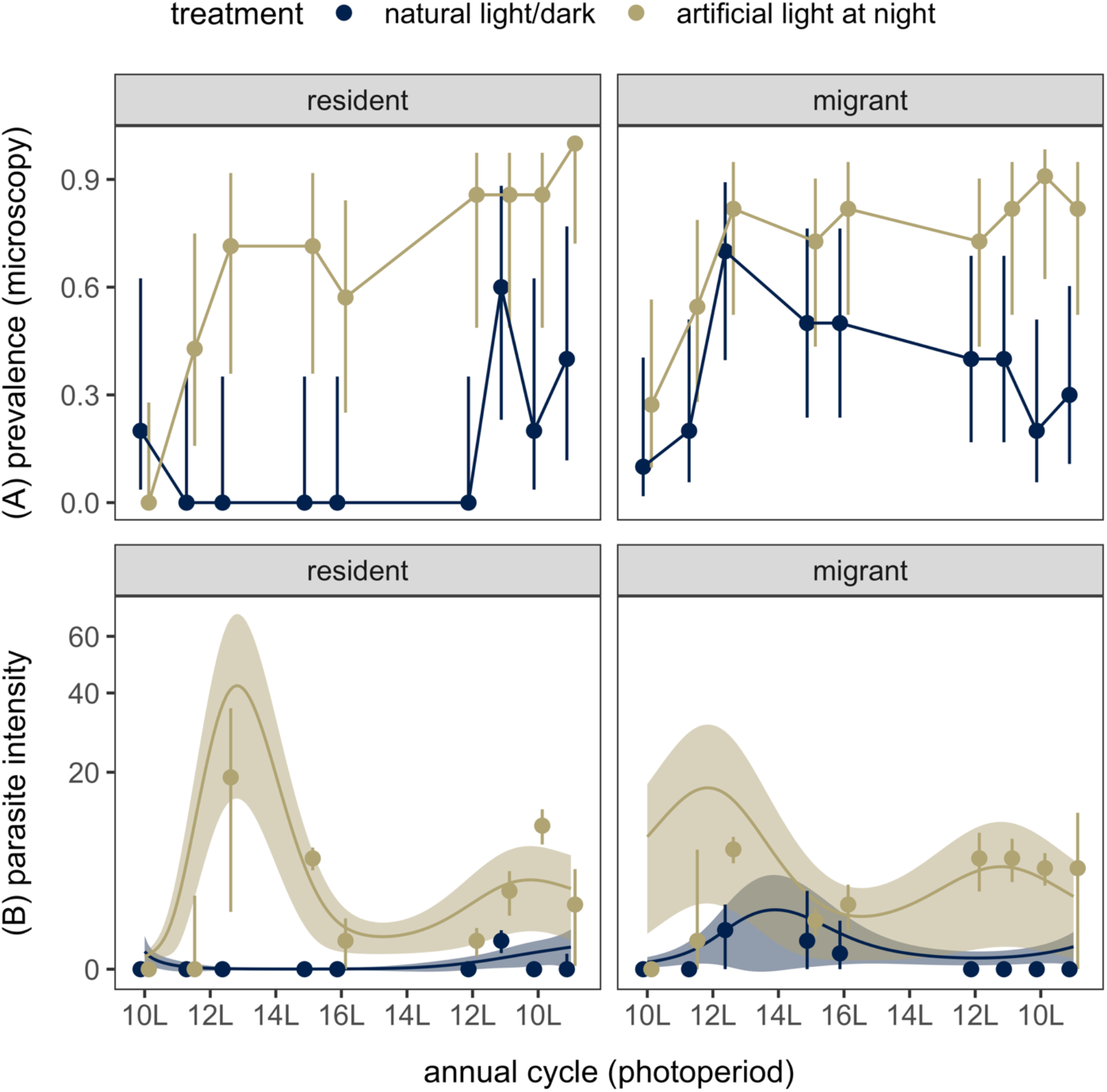
Effects of ALAN on haemosporidian infection dynamics across the simulated annual cycle (i.e., photoperiod, L) of resident and migrant dark-eyed juncos. (A) Prevalence of microscopy-detectable infection is displayed with points and 95% confidence intervals. (B) Parasite intensity is shown as medians with jackknife estimates of standard error. Lines and shaded bands show the fitted values and 95% confidence interval from the GAMM. The vertical axes are displayed with a modulus transformation to accommodate lower bounds at zero. All points are jittered to reduce overlap.

Our GAMMs explained up to 63% of the deviance in haemosporidian intensity and revealed marked effects of ALAN on seasonal parasitemia (Fig. 2B). Haemosporidian intensity in juncos was higher under ALAN compared to natural LD conditions, although this effect was greater in residents when compared to migrants (Table 3). For all ALAN birds, parasitemia peaked around 12L, corresponding to spring, followed by a secondary but smaller parasite intensity peak in photoperiods corresponding to autumn (i.e., 12–10L). Control residents had relatively minor seasonal changes in parasitemia, whereas control migrants also increased haemosporidian intensity in photoperiods corresponding to both spring and fall (Fig. 2B).

**Table 3.**
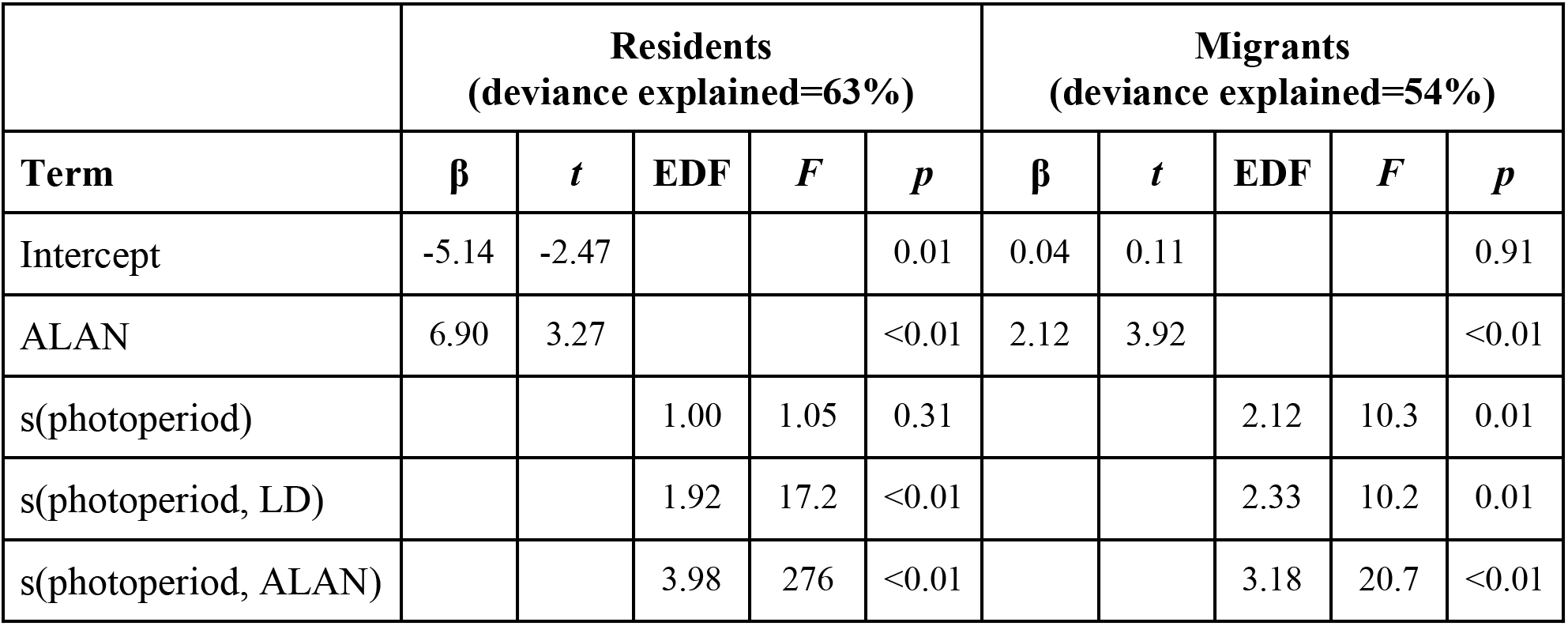
Effects of ALAN treatment, photoperiod, and their interaction on parasite intensity. GAMMs were stratified by migratory strategy and fit using restricted maximum likelihood. Fixed effects are presented with model coefficients (categorical) or the estimated degrees of freedom (EDF) and test statistics. Deviance explained (%) is presented for both GAMMs.

### Cross-correlation between immunity and infection

ALAN affected the cross-correlation between cellular immunity measures and haemosporidian intensity across our experiment, with our GAMMs explaining up to 84% of the deviance in cross-correlation coefficients (Tables S4 and S5). For control residents, correlations between total leukocytes and parasitemia showed strong nonlinearity (*F*=5.43, *p*<0.01) and were most positive near the zero-photoperiod lag, whereas HL ratios had weak effects on parasitemia (*F*=0.20, *p*=0.25; Fig. 3). However, ALAN removed this synchrony, resulting in positive correlations between total leukocytes and haemosporidian intensity across all lags and a non-significant trend (*F*=0, *p*=0.13). For migrants, LD birds instead showed negative correlations around the zero lag (i.e., when the time series were in phase) between total leukocytes and parasitemia (*F*=18.9, *p*<0.01) and positive correlations between HL ratios and parasitemia (*F*=24.6, *p*<0.01; Fig. 3). Similar to residents, ALAN weakened migrant immunity–intensity relationships for total leukocytes (*F*=2.63, p=0.02) and HL ratios (*F*=7.06, *p*<0.01; Fig. 3).

**Figure 3.**
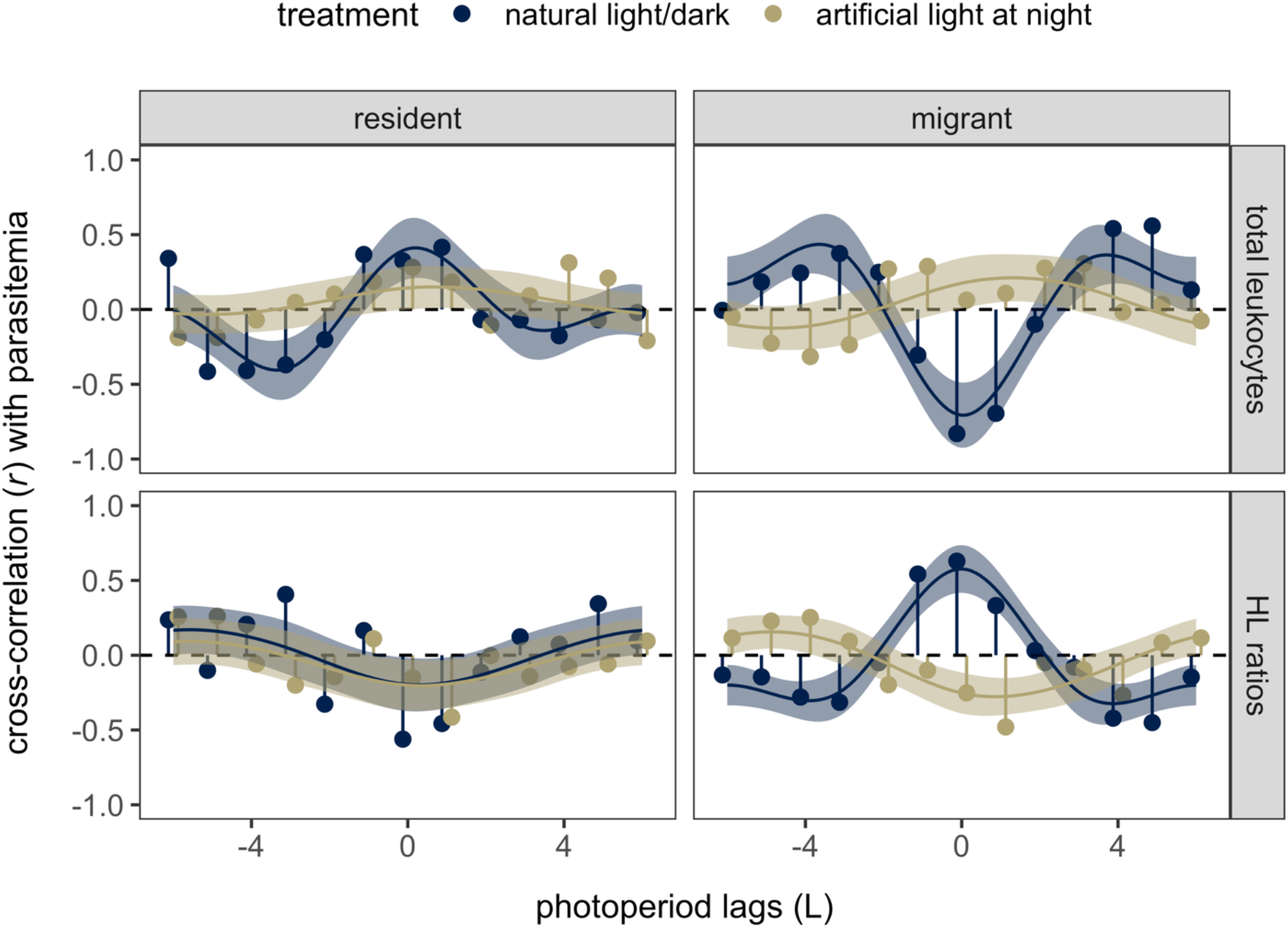
Cross-correlation between immune measures (total leukocytes and HL ratios) and haemosporidian intensity throughout the duration of the experiment. Points display correlation coefficients (*r*) at each photoperiod lag and migratory strategy and are colored by treatment. Lines and shaded bands display the fitted values and 95% confidence interval from each GAM.

## Discussion

Understanding how anthropogenic habitats affect host–parasite interactions is important to forecast how these environments will shape infection risks for wildlife and humans. Here, we experimentally showed that ecologically relevant levels of ALAN, a pervasive component of suburban and urban habitats, increased total leukocytes and haemosporidian intensity in resident and migratory songbirds. Treatment birds had pronounced parasitemia peaks in the spring and fall that were largely absent in control birds, indicating that ALAN caused relapse of chronic infection during energetically costly life history stages. Further, although our immunological and parasitological time series were in phase for control birds, ALAN desynchronized leukocyte profiles and haemosporidian intensity, which could suggest general immune dysregulation.

Because our study occurred in an indoor aviary and PCR confirmed many birds had chronic infections, elevated total leukocytes and haemosporidian intensity in birds exposed to ALAN reflect altered within-host infection dynamics rather than acute infections resulting from vector exposure [32]. In some contexts, ALAN can be immunosuppressive [8,9]; however, the elevated leukocyte counts observed here more likely indicate inflammation in response to haemosporidians [46]. In mice, ALAN can induce an exaggerated inflammatory response driven by proinflammatory cytokines, excessive production of which causes immunopathology and can limit control of infection [10,11]. Similarly, zebra finches (*Taeniopygia guttata*) exposed to ALAN also displayed neuroinflammation with these cytokines [47]. The mechanisms by which the avian immune system controls haemosporidians remains unclear [48], but adaptive responses are likely also important [49,50]. Related to a possibly exaggerated inflammatory response, total leukocytes were positively associated with parasite intensity regardless of time lag, whereas control birds showed more acute synchrony with parasitemia. Notably, we observed these immunological effects at a lower light intensity (2.5 lux) than similar studies (e.g., 5 to 400 lux) [8–11,13,47], suggesting that chronic exposure to even dim ALAN could cause immune dysregulation. However, further quantification of immune gene expression would help identify the specific pathways by which ALAN affects reactivation of haemosporidian infection [51].

Although our results agree with the prediction that ALAN would promote inflammation, we observed generally weak effects on HL ratios. HL ratios can signal elevated glucocorticoids in response to stressors [33], although increases occur more slowly than plasma corticosterone and can be caused by not only intense stressors but also more mild and chronic challenges [52]. Our results thus suggest chronic stimulation of the hypothalamic–pituitary–adrenal axis may not be the underlying mechanism linking ALAN with parasite relapse. This finding is similar to other work on ALAN and within-host processes (e.g., infectious periods for West Nile virus in house sparrows), which suggested relationships between ALAN and increased viremia were not mediated by corticosterone [13]. Alternatively, relationships between ALAN, immunity, and parasitemia could be mediated by other seasonally varying hormones such as testosterone or melatonin [53]. In particular, melatonin is secreted for a longer duration in short days and can facilitate immunostimulation [54]. Additionally, fluctuations in host melatonin can influence life cycle periodicity of some haemosporidians [25]. Suppression of melatonin with ALAN could explain seasonal relapse during periods in which immunity would otherwise be enhanced [55]. Future work could experimentally assess the hormonal basis of ALAN-induced parasite relapse.

Residents and migrants generally responded in a similar fashion to ALAN, with elevated leukocytes and increased parasitemia in spring and fall. However, both control and treatment migrants displayed reactivation of chronic infections in spring, likely signaling physiological changes in preparation for migration combined with those for reproduction [29,30]. In contrast, whereas residents exposed to ALAN also experienced dramatic spring relapse, control residents showed only weak increases in parasitemia in late autumn. These patterns suggest that effects of ALAN on parasite relapse may be greater during periods of reproduction rather than preparation for migration, especially as complementary work demonstrated that ALAN also advanced timing of junco reproduction [37]. Mounting an immune response against haemosporidians can be costly, which could drive weaker parasite control during energetic trade-offs with reproductive activity [48,56]. However, our study only assessed transitions into migratory preparedness and not long-distance migration itself, which could impose stronger effects on parasite control and therefore on cycles of latency and reactivation [57,58]. Lastly, fall increases in parasitemia for both residents and migrants under ALAN could again signal melatonin suppression [53,55].

Although the mechanisms linking ALAN with parasite relapse require further study, particularly for non-migratory hosts, our findings also have implications for the population-level consequences of infection. For haemosporidian parasites, fitness costs for birds can be highly variable [25]. In some cases, haemosporidians have caused dramatic population declines in hosts such as some Hawaiian honeycreepers (e.g., *Vestiaria coccinea*) and possibly house sparrows (*Passer domesticus*) in Europe [59,60]. Relapse of chronic infections under ALAN in anthropogenic habitats could accordingly either facilitate infection-related population declines in less tolerant avian hosts or increase parasitemia in tolerant species that instead serve as sources of infection for generalist vectors, thereby increasing transmission in the local avian community. To reduce such negative impacts, alternative lighting schemes in suburban and urban environments, such as warmer light spectra, minimizing upwards-facing lighting structures, and restricting use of broad-spectrum light during demanding stages of avian annual cycles (e.g., migration), warrant further investigation as ecologically friendly interventions [61–63]. Interdisciplinary collaborations between physiologists and urban planners could enable testing such interventions in field experiments and facilitate their adoption into these landscapes.

More generally, our study highlights reactivation of chronic or latent infection as a novel mechanism by which urbanization can affect infectious disease dynamics. Examining how ALAN affects cycles of latency and reactivation in other host and parasite systems, such as *Borrelia burgdorferi* and arboviruses in songbirds [64,65] or henipaviruses and herpesviruses in bats [66,67], could inform diseases risks relevant for wildlife conservation, domestic animal health, and human health. Such work could be especially informative for synanthropic wildlife and migratory species that are increasingly becoming sedentary in anthropogenic habitats [68].

## Author contributions

DS, DJB, and EDK designed the study; DS, DJB, JDM, KMT, and SW collected samples; QP analyzed blood smears; KMT conducted PCR; and DJB analyzed data and wrote the manuscript. All authors contributed to revisions.

## Acknowledgements

We thank members of the Ketterson lab and two reviewers for helpful feedback.

## Ethics

All procedures were approved by the Indiana University Institutional Animal Care and Use Committee (18-030) and conducted under permits issued by the Virginia Department of Game and Inland Fisheries (052971) and U.S. Fish and Wildlife Service (20261).

## Funding

This work was supported by the Environmental Resilience Institute, funded by Indiana University’s Prepared for Environmental Change Grand Challenge Initiative. DJB was supported by an appointment to the Intelligence Community Postdoctoral Research Fellowship Program, administered by Oak Ridge Institute for Science and Education through an interagency agreement between the U.S. Department of Energy and Office of the Director of National Intelligence.

## Data availability

Data are available in the Dryad Digital Repository: https://datadryad.org/stash/share/IxJYbB61kJJifIIXuEC72ZcIPEpblW6TjIII6sQ8rQM

## Notes

### Competing Interest Statement

The authors have declared no competing interest.

