## Supplemental Material for "Artificial light at night amplifies seasonal relapse of haemosporidian parasites in a widespread songbird"

Table S1. ANOVA table for generalized linear mixed models (using the *lme4* package) for the fixed effects of migratory strategy, treatment, and their interaction on junco response variables. Models include a random effect of individuals, and error structures are identical to the GAMMs (with the exception of haemosporidian intensity, which used a negative binomial distribution).

| **Response** | **Predictor** | $\chi^2$ | ***p*** |
|---|---|---|---|
| Total leukocytes | Treatment | 5.99 | 0.01 |
| | Migratory | 0.004 | 0.95 |
| | Interaction | 0.003 | 0.96 |
| HL ratios | Treatment | 0.38 | 0.54 |
| | Migratory | 0.26 | 0.61 |
| | Interaction | 0.03 | 0.87 |
| Parasite intensity | Treatment | 34.19 | <0.001 |
| | Migratory | 1.94 | 0.16 |
| | Interaction | 2.17 | 0.14 |

Table S2. Effects of ALAN treatment, photoperiod, and their interaction on HL ratios. GAMMs were stratified by migratory strategy and fit using restricted maximum likelihood. Fixed effects are presented with model coefficients (categorical) or the estimated degrees of freedom (EDF) and test statistics. Deviance explained (%) is presented for both GAMMs.

| | **Residents (deviance explained=44%)** | | | | | **Migrants (deviance explained=45%)** | | | | |
|---|---|---|---|---|---|---|---|---|---|---|
| **Term** | **β** | ***t*** | **EDF** | ***F*** | ***p*** | **β** | ***t*** | **EDF** | ***F*** | ***p*** |
| Intercept | -1.69 | -7.57 | | | <0.01 | -1.90 | -12.1 | | | <0.01 |
| ALAN | -0.28 | -0.94 | | | 0.35 | -0.31 | -1.43 | | | 0.15 |
| s(photoperiod) | | | 1.00 | 1.03 | 0.31 | | | 3.22 | 8.47 | <0.01 |
| s(photoperiod, LD) | | | 2.21 | 1.01 | 0.27 | | | 1.00 | 0.00 | 1.00 |
| s(photoperiod, ALAN) | | | 1.62 | 1.57 | 0.19 | | | 0.00 | >10 | <0.01 |

Table S3. Effects of ALAN treatment, migratory strategy, and their interaction on the likelihood of detecting haemosporidians by PCR at the start of the experiment (9L, 2/2019). Shown are test statistics and $p$ values from a GLM with binomial errors and a logit link.

| **Predictor** | $\chi^2$ | $p$ |
|---|---|---|
| Treatment | 0.63 | 0.43 |
| Migratory | 0.06 | 0.80 |
| Interaction | 0.54 | 0.46 |

Table S4. Effects of ALAN treatment, photoperiod lag, and their interaction on correlation coefficients between total leukocytes ($\log_{10}$) and log(x+1)-transformed haemosporidian intensity. GAMMs were stratified by migratory strategy and fit using restricted maximum likelihood. Fixed effects are presented with model coefficients (categorical) or the estimated degrees of freedom (EDF) and test statistics. Deviance explained (%) is presented for both GAMMs.

| | **Residents (deviance explained=64%)** | | | | | **Migrants (deviance explained=78%)** | | | | |
|---|---|---|---|---|---|---|---|---|---|---|
| **Term** | **β** | ***t*** | **EDF** | ***F*** | ***p*** | **β** | ***t*** | **EDF** | ***F*** | ***p*** |
| Intercept | -0.03 | -0.58 | | | 0.57 | 0.04 | 0.86 | | | 0.40 |
| ALAN | 0.08 | 0.86 | | | 0.25 | -0.01 | -0.13 | | | 0.90 |
| s(lag) | | | 1.32 | 1.14 | 0.09 | | | 0.00 | 0.00 | 0.04 |
| s(lag, LD) | | | 2.76 | 5.43 | <0.01 | | | 2.87 | 18.9 | <0.01 |
| s(lag, ALAN) | | | 0.00 | 0.00 | 0.13 | | | 1.74 | 2.63 | 0.02 |

Table S5. Effects of ALAN treatment, photoperiod lag, and their interaction on correlation coefficients between HL ratios (square-root) and log(x+1)-transformed haemosporidian intensity. GAMMs were stratified by migratory strategy and fit using restricted maximum likelihood. Fixed effects are presented with model coefficients (categorical) or the estimated degrees of freedom (EDF) and test statistics. Deviance explained (%) is presented for both GAMMs.

| | **Residents (deviance explained=38%)** | | | | | **Migrants (deviance explained=84%)** | | | | |
|---|---|---|---|---|---|---|---|---|---|---|
| **Term** | **β** | ***t*** | **EDF** | ***F*** | ***p*** | **β** | ***t*** | **EDF** | ***F*** | ***p*** |
| Intercept | 0.01 | 0.12 | | | 0.90 | -0.04 | -1.04 | | | 0.31 |
| ALAN | -0.05 | -0.58 | | | 0.57 | -0.01 | -0.11 | | | 0.92 |
| s(lag) | | | 1.72 | 2.45 | 0.01 | | | 0.00 | 0.00 | 0.03 |
| s(lag, LD) | | | 0.48 | 0.20 | 0.25 | | | 2.87 | 24.6 | <0.01 |
| s(lag, ALAN) | | | 0.00 | 0.00 | 0.95 | | | 2.16 | 7.06 | <0.01 |

Figure S1. Prevalence of PCR-detectable haemosporidian infections in blood for resident and migrant juncos under control and treatment conditions at the start of the experiment (9L, $n$ = 33). Lines display 95% confidence intervals. PCR reactions were run in 10 µL volumes, including 2 µL of DNA, 5 µL of Promega PCR master mix, and 0.8 µM of each primer (343F and 496R).

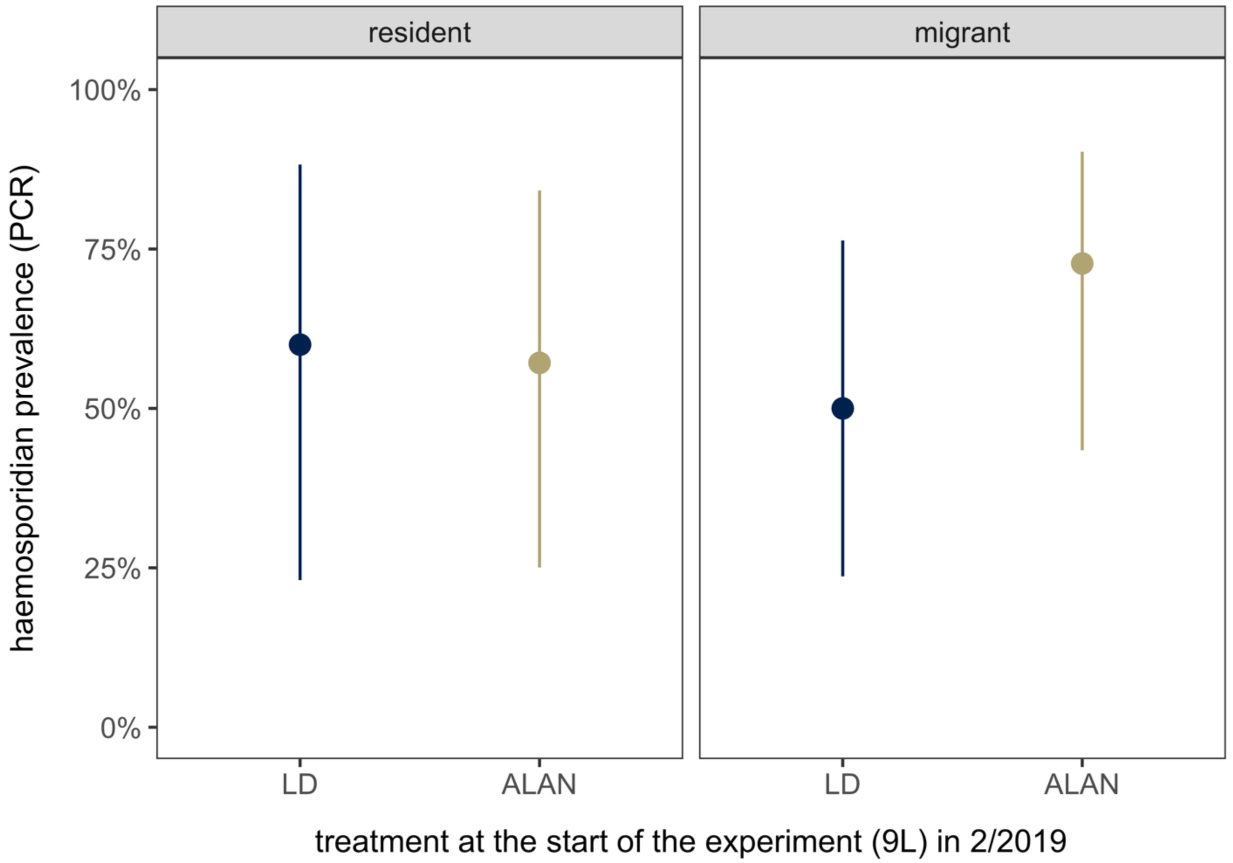

Figure S2. Concordance between haemosporidian PCR at the start of the experiment (9L) and parasite intensity (microscopy) at the subsequent 10L timepoint. Most 9L PCR-negative birds remained microscopy-negative at 10L (92%). The vertical axis is displayed with a modulus transformation. Points are shaped by binary microscopy detection and are colored by treatment.

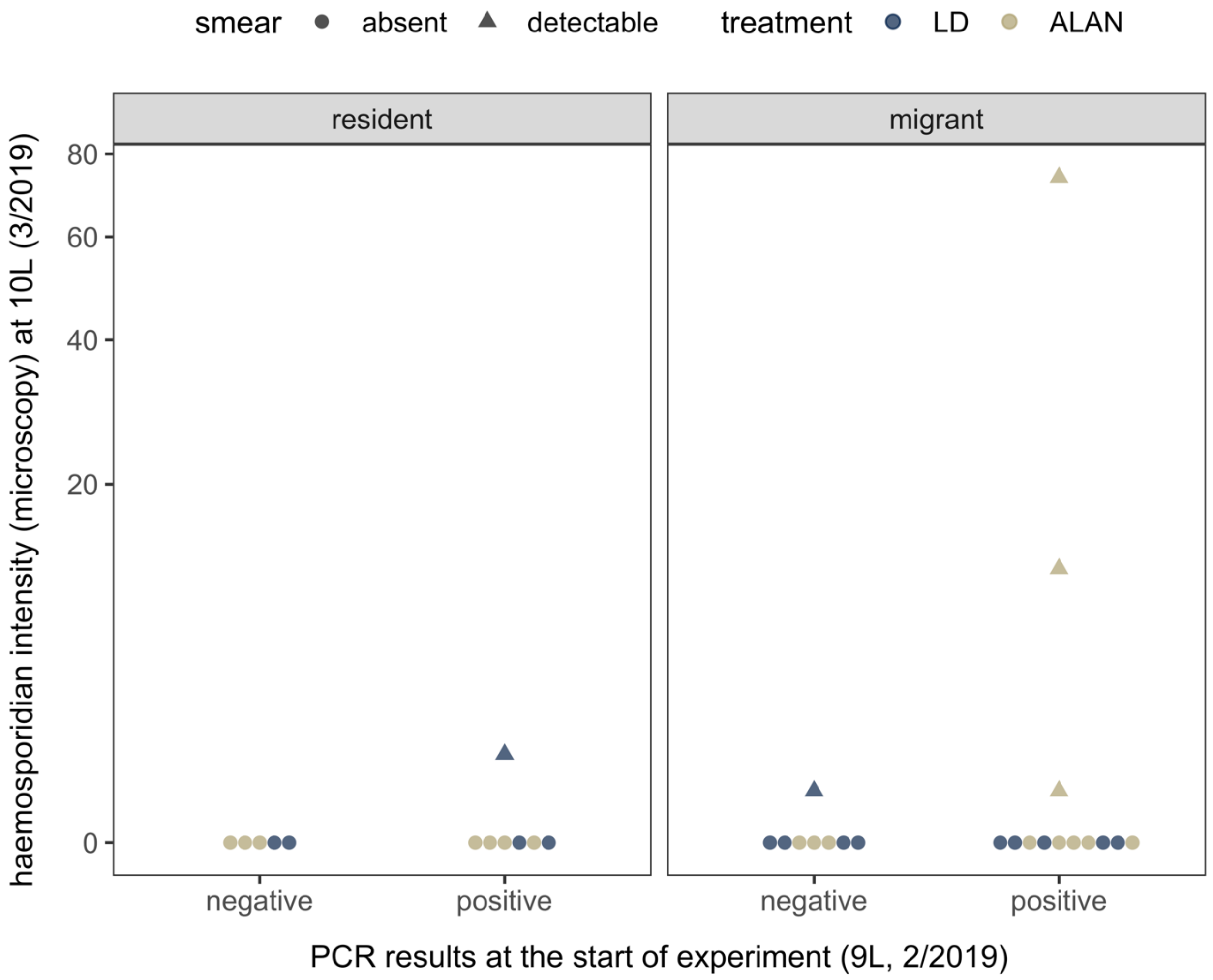
